# Melatonin attenuates kidney injury by alleviating lysosomal damage in diabetic kidney disease

**DOI:** 10.1101/2025.01.06.631433

**Authors:** Jiaqi Chen, Shuting Zhang, Xiaoqin Ma, Aomiao Chen, Yichuan Wu, Geningyue Wang, Qian Zhang, Yaoming Xue, Yijie Jia, Zongji Zheng

**Affiliations:** Department of Endocrinology and Metabolism, Nanfang Hospital, Southern Medical University, Guangzhou, China; Department of Endocrinology, Guangdong Provincial People’s Hospital (Guangdong Academy of Medical Sciences), Southern Medical University, Guangzhou, China; De Feng Academy, Southern Medical University, Guangzhou, China

**Keywords:** diabetic kidney disease, melatonin, lysosome, TFEB, tubular epithelial cell

## Abstract

Proteinuria-induced damage to renal tubular epithelial cells is one of the main causes of diabetic kidney disease (DKD), and the clearance of overloaded albumin by lysosomes is crucial for maintaining the homeostasis of renal tubular epithelial cells. Therefore, lysosomal damage is closely related to the pathogenesis of DKD, but effective prevention and treatment measures are still lacking. Melatonin (MLT) is secreted by the pineal gland and can not only regulate circadian rhythms but also maintain lysosomal homeostasis. In this study, we demonstrate the presence of significant lysosomal damage in renal tubules of DKD patients, which causes autophagy impairment and a concomitant oxidative stress imbalance; however, MLT can upregulate transcription factor EB (TFEB) to improve lysosomal damage and restore the biosynthesis in this organelle. Mechanistically, MLT may protect lysosomes via the upregulation of TFEB and the miR-205-5p‒LRP-1 pathway in renal tubules, thus improving autophagy dysfunction and oxidative imbalance in DKD.

## INTRODUCTION

Diabetic kidney disease (DKD) is a microvascular disease caused by diabetes that is associated with significant mortality and disability [1]. Regrettably, few effective DKD treatments currently exist [2]. Thus, the discovery of novel pathophysiological mechanisms and the formulation of novel treatment approaches for DKD are needed. Urinary albumin is not only the most characteristic clinical manifestation of DKD but is also an independent risk factor for its progression [3, 4]. Previous studies have shown that prolonged exposure to an excessive amount of albumin can result in the overloading of albumin in renal tubular epithelial cells, which induces inflammatory responses, oxidative stress, and apoptosis[5, 6]. In addition, increasing evidence suggests that autophagy can protect renal tubular epithelial cells from damage in various kidney diseases, including DKD [7, 8].

As the main degradation system in cells, autophagy plays an important role in adapting to environmental changes, maintaining internal homeostasis, responding to stress and maintaining cellular function [9]. The degradative capacity of autophagy originates in lysosomes; therefore, lysosomes occupy a position at the core of the autophagic process. Although lysosomes are essential for autophagy, lysosomes are mostly neglected in autophagy studies, and thus the means needed to alleviate lysosomal damage are very limited [10]. Several studies have shown that, in DKD, albumin and glucose overload can cause damage to lysosomes in renal tubular epithelial cells and can subsequently promote DKD occurrence and development [11–13]. Liu *et al*. reported that lysosomal dysfunction throughout the course of DKD represents the primary mechanism underlying the protective potential of autophagy [14]. Moreover, lysosomal repair can activate autophagy and improve kidney homeostasis [15, 16]. Therefore, inhibition of lysosomal damage and depletion can prevent peroxidation, maintain homeostasis in renal tubular epithelial cells, and thereby improve DKD. Nevertheless, the mechanism of lysosomal damage and repair in renal tubules in DKD is still unclear.

MLT was initially believed to be a hormone secreted by the pineal gland that regulates circadian rhythms [17]. With increasing research, scientists have discovered that MLT also has powerful functions in antioxidant stress, elimination of inflammation and induction of autophagy, and thus MLT plays an important protective role in multiple diseases [18, 19]. Recently, the mechanisms of DKD have been partially revealed.

Several studies have shown that MLT intervention can prevent and significantly ameliorate the damage caused by oxidative stress, apoptosis and inflammation in the kidneys of diabetic rodents [20–22]. In vitro studies have also shown that MLT can exert anti-inflammatory and antioxidant effects, thus alleviating kidney podocyte, endothelial cell and mesangial cell damage in the diabetic state [23, 24]. However, these studies focused mainly on the glomeruli and largely ignored the renal tubules, which also play important roles in DKD. Recent studies have shown that MLT may delay the progression of DKD by increasing mitophagy in the proximal renal tubules [25]. Unfortunately, the relationships among MLT, autophagic core lysosomes and renal tubules are still not well studied in DKD. Moreover, Liu *et al*. reported that the oral administration of MLT increased TFEB levels, which increased lysosomal function; this in turn restored autophagic flux to mitigate doxorubicin-induced cardiomyopathy [26]. In an animal model of sciatic nerve injury, MLT treatment increased the number of lysosomes and ultimately promoted nerve regeneration[27]. Although no studies have shown that MLT delays the progression of DKD by alleviating lysosomal damage, the results of the aforementioned studies suggest that MLT has potential as a prospective pharmaceutical option for the treatment of DKD.

In this study, we explored the potential of MLT to ameliorate lysosomal damage in renal tubular epithelial cells, restore autophagic flux, mitigate oxidative stress, and consequently slow the progression of DKD.

## MATERIALS AND METHODS

### Cell Culture and Experimental Procedures

Experiments were conducted using human proximal tubular HK-2 cells, which were obtained from the Chinese Academy of Sciences in Shanghai. The cells were cultured in DMEM/F12 media supplemented with 10% fetal bovine serum (Gibco, USA) at 37°C in an incubator containing 5% carbon dioxide.

For treatment with bovine serum albumin (BSA, Sigma, V900933), HK-2 cells were cultured in medium supplemented with 2% FBS for 24 hours after which they were subsequently exposed to 10 mg/mL BSA for 48 hours. To evaluate the regulatory role of MLT in countering BSA-induced injury, 100 µM MLT (MCE, HY-B0075) was administered at the time of BSA stimulation.

For the transfection of HK-2 cells with a mimic or an inhibitor of miR-205-5p (RiboBio, Guangzhou, China), a miR-205-5p-5p mimic (50 nM) or its corresponding negative control was used; alternatively, a miR-205-5p inhibitor (100 nM) or its negative control was used. Transfections were performed for 24 hours using a Lipofectamine 3000 Kit (Invitrogen, L3000015). HK-2 cells were seeded in a six-well plate (GeneCopoeia, Guangzhou, China) and transfected with 2.5 μg of the pEZ-Lv201-LRP-1 or pEZ-Lv201-TFEB plasmid to overexpress LRP-1 or TFEB, respectively. The transfection duration was 48 hours, according to the manufacturer’s guidelines. Additionally, to investigate the influence of MLT on lysosome depletion via TFEB, we transfected HK-2 cells with 50 nM TFEB siRNA (RiboBio, Guangzhou, China) or negative control siRNA for 48 hours. All in vitro experiments were conducted in triplicate, and each treatment was performed in duplicate.

### Animal Experiments

We obtained db/db mice and nondiabetic control db/m mice (male; 8 weeks old) from the Nanjing Biomedical Research Institute (China). These mice were raised in a specific pathogen-free environment and fed a standard diet. A subset of these mice (five db/m and five db/db) was sacrificed when they reached 24 weeks of age.

To determine the effect of MLT on DKD, ten db/db mice (20 weeks old) were equally divided into two groups: db/db+H_2_O control (distilled water, gavage) and db/db+MLT (MCE, HY-B0075, 20 mg/kg, gavage). We continuously treated the mice with MLT 3 times/week for 4 weeks. When the mice reached 24 weeks of age, to euthanize the animals, we administered intraperitoneal injections of pentobarbital sodium (100 mg/kg).

To reveal the protective effect of miR-205 on DKD, ten db/db mice (20 weeks old) were equally divided into two groups: db/db+agomiR-negative control (agomiR-NC) and db/db+miR-205 agonist (RiboBio, Guangzhou, China; 2.5 mg/kg, tail vein injection). We continuously treated the mice twice per week for one month. When the mice reached 24 weeks of age, the animals were euthanized by intraperitoneal injections of pentobarbital sodium (100 mg/kg). Approval for all animal experiments was granted by Southern Medical University (Approval Number: L2018022).

### Reverse Transcription-Polymerase Chain Reaction (RT-PCR)

Total RNA was isolated from HK-2 cells and mouse renal cortex using TRIzol (Invitrogen, USA). DNA was synthesized with a test kit (11141ES60, Yeasen, China) for mRNA analysis and a Mir-X™ miRNA First-Strand Synthesis Kit (KR211-02 and FP411-02, TIANGEN, China) for miRNA analysis. The primers used were obtained from Tiangen Biochemical Technology (Beijing) Co., Ltd.

SYBR Green Master Mix (11202ES08, Yeasen, China) was used for qPCR. 18S expression was used for normalization of the mRNA expression level, whereas snRNA U6 expression was used for normalization of the miRNA expression level according to the 2^-ΔΔCt method. The detailed primer sequences (Generay, Shanghai, China) are provided in Supplementary Table 1.

### Western blot (WB) Analysis

We used RIPA lysis buffer (FD008, Fdbio Science, China) supplemented with protease inhibitors and phosphatase inhibitors (78,440, Thermo Fisher Scientific) to extract total protein from both the mouse renal cortex and HK-2 cells. Gel electrophoresis was performed to separate proteins of different molecular weights, which were then transferred to a nitrocellulose membrane (NC membrane; 66485, PALL, China). Antibodies against HO 1 (WL02400, Wanleibio, China), NRF2 (WL02135, Wanleibio, China), p62 (WL02385, Wanleibio, China), LC3 (14600-1-AP, Proteintech, China), GAPDH (60004-1-Ig, Proteintech, China), LRP-1 (WL003211, Wanleibio, China), and TFEB (13372-1-AP, Proteintech, China) were used. We captured images via an enhanced chemiluminescence-based imaging detection system (GelView 6000Pro, Guangzhou, China).

### Reactive Oxygen Species (ROS) Detection in vivo and in vitro

Reactive oxygen species (ROS) were measured via H2DCFDA (MedChemExpress, China) in vitro and in vivo. HK-2 cells and frozen kidney sections were exposed to the dye (10 µM) for 30 minutes at 25°C in the dark. Images were obtained with a fluorescence microscope (Imager D2, Zeiss, Germany).

### Measurement of the Mitochondrial Membrane Potential

A kit containing the fluorescent sensor JC-1 was used to rapidly identify changes in the mitochondrial membrane potential (MedChemExpress, USA). HK-2 cells and frozen kidney sections were incubated with the dye (200 µM) at 25°C in the dark. Images were obtained with a fluorescence microscope (Imager D2, Zeiss, Germany).

### LysoTracker Assays

Kidney tissue sections and cells were exposed to LysoTracker Red dye (50 nM) and incubated at 37°C in the dark. Images were obtained with a fluorescence microscope (Imager D2, Zeiss, Germany).

### Superoxide Dismutase (SOD) Activity Assessment

To assess SOD activity, we used a Total SOD Activity Detection Kit (Beyotime, S0101S).

### Immunofluorescence Staining

For IF staining, we initially fixed cell climbing slides or frozen kidney tissue sections in 4% paraformaldehyde. Triton X-100 was used for permeabilization after which blocking was performed with 5% BSA. Next, cells and tissues were incubated overnight with the following antibodies at a 1:150 dilution: anti-HO1, anti-NRF2, anti-LRP-1 (WL03211, Wanleibio, China), anti-TFEB (13372-1-AP, Proteintech, China), anti-LAMP1 (WL02419, Wanleibio, China), anti-GAL3 (60207-1, Proteintech, China), and anti-LTL (FL-1321-2, Vector, USA). LTL was used as a marker of the proximal renal tubules in the mouse kidney to specifically locate and observe the expression of GAL3, LAMP1, and TFEB at these locations. After the primary antibody was removed, the cells and tissues were incubated with secondary antibodies (E032410 and E032420-01, EarthOX) in the dark for 1 hour. We captured images with a fluorescence microscope (Imager D2, Zeiss, Germany).

### Double Luciferase Reporter Assay

HEK293T cells were used for the dual-luciferase reporter assay. The sequences of the WT and mutant LRP-1 3’-UTR plasmids are provided in Supplementary Table 2. The sequences of the WT and mutant miR-205-5p promoter plasmids are listed in Supplementary Table 3.

## RESULTS

### Albumin induces lysosomal damage in renal tubular epithelial cells and DKD model animals

The oxidative stress imbalance and autophagy arrest caused by albumin overload and lysosomal injury are important causes of the destabilization of renal tubular epithelial cells, which may be closely related to the pathogenesis of DKD. Therefore, we first examined oxidative stress in the renal cortex of mice. Compared with that in db/m mice (the control group), the level of the antioxidant enzyme SOD in db/db mice was significantly lower (Figure 1A), whereas the ROS level was significantly higher (Figure S1), which resulted in a significant decrease in the mitochondrial membrane potential (Figure S2). Moreover, immunofluorescence staining (Figure S3A, B) and WB (Figure 1B) revealed that the levels of the antioxidative stress-related proteins NRF2 and HO1 were also significantly lower in db/db mice than in db/m mice. Fluorescence staining revealed significant lysosomal damage and reduced biosynthesis in db/db mice (Figure 1C-F). Additionally, compared with those in db/m mice, the protein levels of LC3 and p62 in db/db mice were significantly increased (Figure 1G), which indicates that autophagy was inhibited.

**Figure 1.**
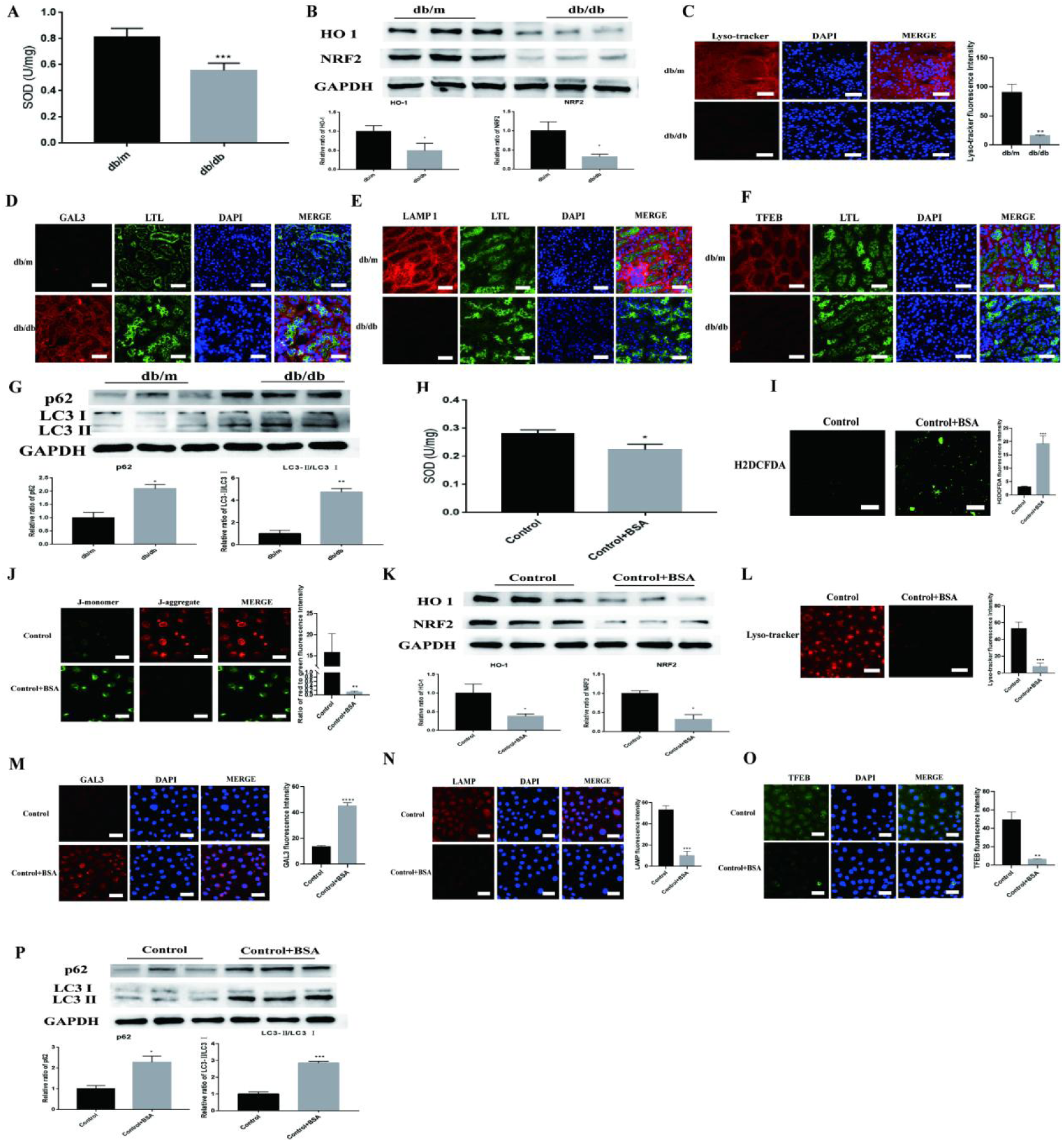
Lysosomal damage and autophagy dysfunction in renal tubular epithelial cells in DKD (A)SOD activity in db/db and control mice (n=5/group). ∗ ∗ ∗ P < 0.001 vs. the control group. (B) Relative HO 1 and NRF2 protein levels in db/db and control mice. ∗ P < 0.05 vs. the control group. (C) Fluorescence intensity of lysosomal acidification in db/db and control mice. ∗ ∗ P < 0.01 vs. the control group. (D-F) Images of GAL3, LAMP1 and TFEB staining in db/db and control mice. (G) Relative p62 and LC3 protein levels in db/db and control mice. ∗ P < 0.05, ∗ ∗ P < 0.01 vs. the control group. (H) SOD activity in BSA-HK-2 cells. ∗ P < 0.05 vs. the control group. (I) Fluorescence intensity of ROS in BSA-HK-2 cells. ∗ ∗ ∗ P < 0.001 vs. the control group. (J) Images of JC-1-stained BSA-HK-2 cells. ∗ ∗ P < 0.01 vs. the control group. (K) Relative HO 1 and NRF2 protein levels in BSA-HK-2 cells. ∗ P < 0.05 vs. the control group. (L) Fluorescence intensity of lysosomal acidification in BSA-HK-2 cells. ∗ ∗ ∗ P < 0.001 vs. the control group. (M-O) Images of GAL3, LAMP1 and TFEB staining in BSA-HK-2 cells. ∗ ∗ P < 0.01, ∗ ∗ ∗ P < 0.001, ∗ ∗ ∗ ∗ P < 0.0001 vs. the control group. (P) Relative p62 and LC3 protein levels in BSA-HK-2 cells. ∗ P < 0.05, ∗ ∗ P < 0.01 (vs. the control group).

We also examined oxidative stress and autophagy in BSA-stimulated HK-2 (BSA-HK-2) cells. The results revealed that BSA-HK-2 cells had significantly reduced SOD levels (Figure 1H) and significantly increased ROS levels (Figure 1I), which caused a significant decrease in the mitochondrial membrane potential (Figure 1J). Immunofluorescence staining (Figure S4A, B) and WB (Figure 1K) were used to assess the expression levels of the antioxidative stress-related proteins NRF2 and HO1 in BSA-HK-2 cells. In addition, fluorescence staining revealed significant lysosomal damage and reduced biosynthesis in BSA-HK-2 cells (Figure 1L-O). The protein levels of LC3 and p62 were also significantly increased (Figure 1P), which suggests the occurrence of autophagy arrest in BSA-HK-2 cells. These results indicate that the oxidative stress imbalance and autophagy arrest in renal tubular epithelial cells caused by albumin overload and lysosomal injury are closely associated with DKD.

### MLT alleviates the lysosomal damage induced by albumin in renal tubular epithelial cells

MLT alleviates kidney injury in DKD models, but the underlying mechanism is unclear. During DKD, significant lysosomal damage and renal tubular epithelial cell depletion occur. To elucidate the mechanism by which MLT improves DKD, we treated BSA-HK-2 cells with MLT. We first examined oxidative stress and autophagy in BSA-HK-2 cells treated with MLT. Compared with untreated BSA-HK-2 cells, BSA-HK-2 cells treated with MLT presented significantly higher levels of SOD (Figure 2A), whereas ROS levels were lower (Figure 2B), which increased the mitochondrial membrane potential (Figure 2C). Moreover, immunofluorescence staining (Figure 2D, E) and WB (Figure 2F) revealed that the levels of the antioxidative stress-related proteins NRF2 and HO1 were also significantly increased in MLT-treated BSA-HK-2 cells. In addition, fluorescence staining revealed lysosomal damage repair and increased biosynthesis in MLT-treated BSA-HK-2 cells (Figure 2G-J). Moreover, the protein levels of LC3 and p62 were significantly decreased after MLT treatment (Figure 2K), which suggests improved autophagy arrest. These results indicate that MLT can reverse the oxidative stress imbalance and autophagy arrest in renal tubular epithelial cells caused by albumin overload and lysosomal injury.

**Figure 2.**
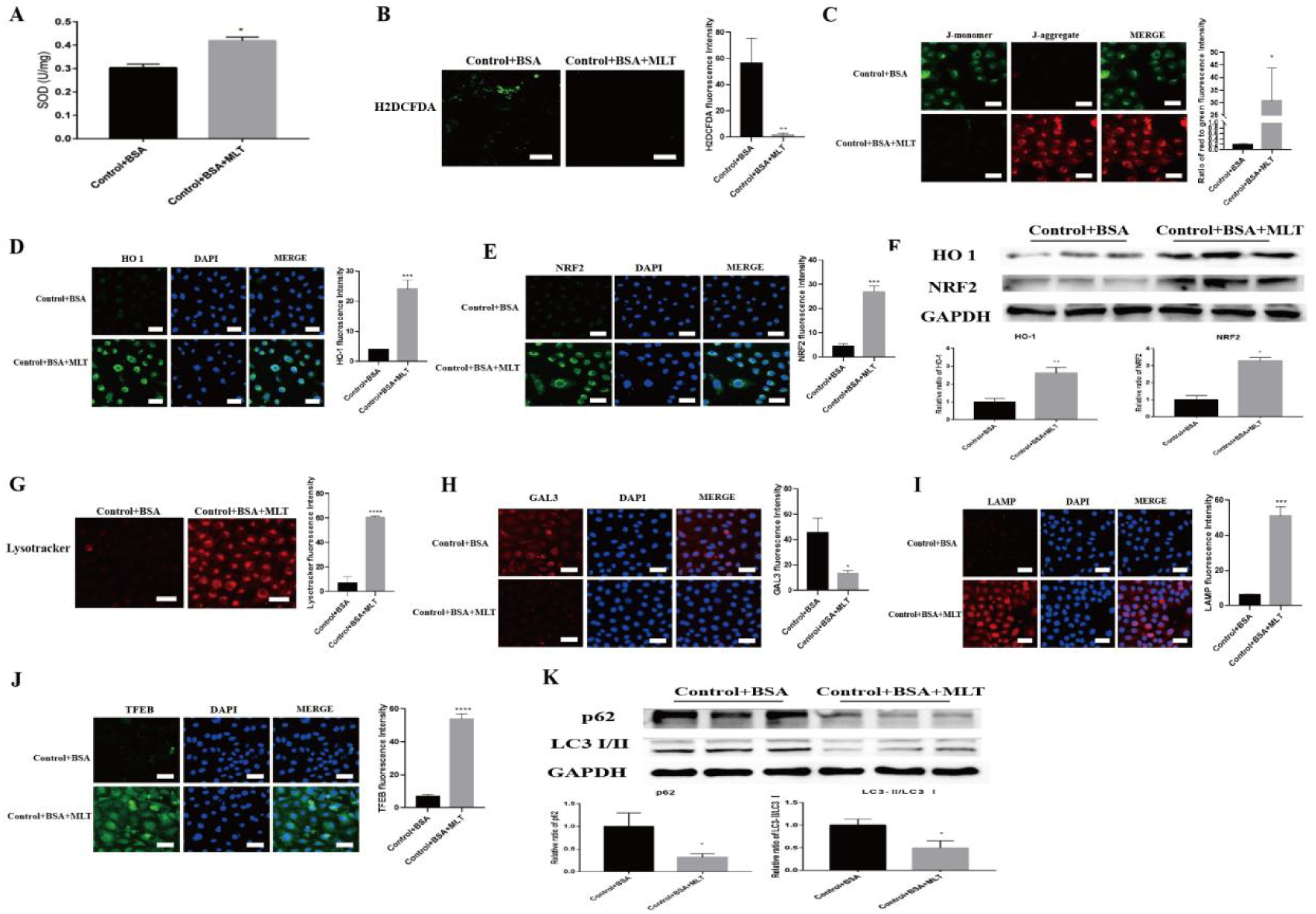
MLT ameliorates lysosomal damage in BSA-HK-2 cells (A) SOD activity in MLT-treated BSA-HK-2 cells. ∗ P < 0.05 vs. the control group. (B) Fluorescence intensity of ROS in MLT-treated BSA-HK-2 cells. ∗ ∗ P < 0.01 vs. the control group. (C) JC-1 staining images of MLT-treated BSA-HK-2 cells. ∗ P < 0.05 vs. the control group. (D, E) Immunofluorescence staining for HO 1 and NRF2 in MLT-treated BSA-HK-2 cells. ∗ ∗ ∗ P < 0.001 vs. the control group. (F) Relative HO 1 and NRF2 protein levels in MLT-treated BSA-HK-2 cells. ∗ P < 0.05 vs. the control group. (G) Fluorescence intensity of lysosomal acidification in MLT-treated BSA-HK-2 cells. ∗ ∗ ∗ ∗ P < 0.0001 vs. the control group. (H-J) Images of GAL3, LAMP1 and TFEB staining in MLT-treated BSA-HK-2 cells. ∗ P < 0.05, ∗ ∗ ∗ P < 0.001, ∗ ∗ ∗ ∗ P < 0.0001 vs. the control group. (K) Relative p62 and LC3 protein levels in MLT-treated BSA-HK-2 cells. ∗ P < 0.05 vs. the control group.

### MLT ameliorates lysosomal damage in BSA-HK-2 cells via miR-205-5p

MiRNAs are closely related to the pathogenesis of DKD, but it is not yet clear whether MLT exerts its effects through miRNAs. Therefore, we next explored the possible mechanism by which MLT inhibits oxidative stress. We screened for miRNAs reported to be associated with renal tubular epithelial cell injury in MLT-treated BSA-HK-2 cells and found that MLT significantly increased the expression of miR-205-5p (Figure 3A), which suggests that miR-205-5p may participate in the mechanism by which MLT protects against DKD. To further confirm that MLT alleviated lysosomal damage in BSA-HK-2 cells through miR-205-5p, we treated BSA-HK-2 cells with MLT and concomitantly suppressed miR-205-5p expression. We subsequently examined oxidative stress, lysosomal damage, and the expression of autophagy-related markers. Compared with BSA-HK-2 cells treated with MLT alone, the inhibition of miR-205-5p expression in MLT-treated BSA-HK-2 cells resulted in a decrease in SOD levels (Figure 3B) and an increase in ROS levels (Figure 3C), which led to a significant decrease in the mitochondrial membrane potential (Figure 3D). Immunofluorescence staining (Figure 3E, F) and WB (Figure 3G) revealed that the levels of the antioxidative stress-related proteins NRF2 and HO1 were also significantly decreased after miR-205-5p knockdown. In addition, fluorescence staining revealed significant lysosomal damage and reduced biosynthesis after miR-205-5p knockdown (Figure 3G-K). The protein levels of LC3 and p62 were also significantly increased (Figure 3L), which suggests the occurrence of autophagy arrest after the inhibition of miR-205-5p. These results indicate that the knockdown of miR-205-5p could block the lysosomal-therapeutic effect of MLT on BSA-HK-2-cells.

**Figure 3.**
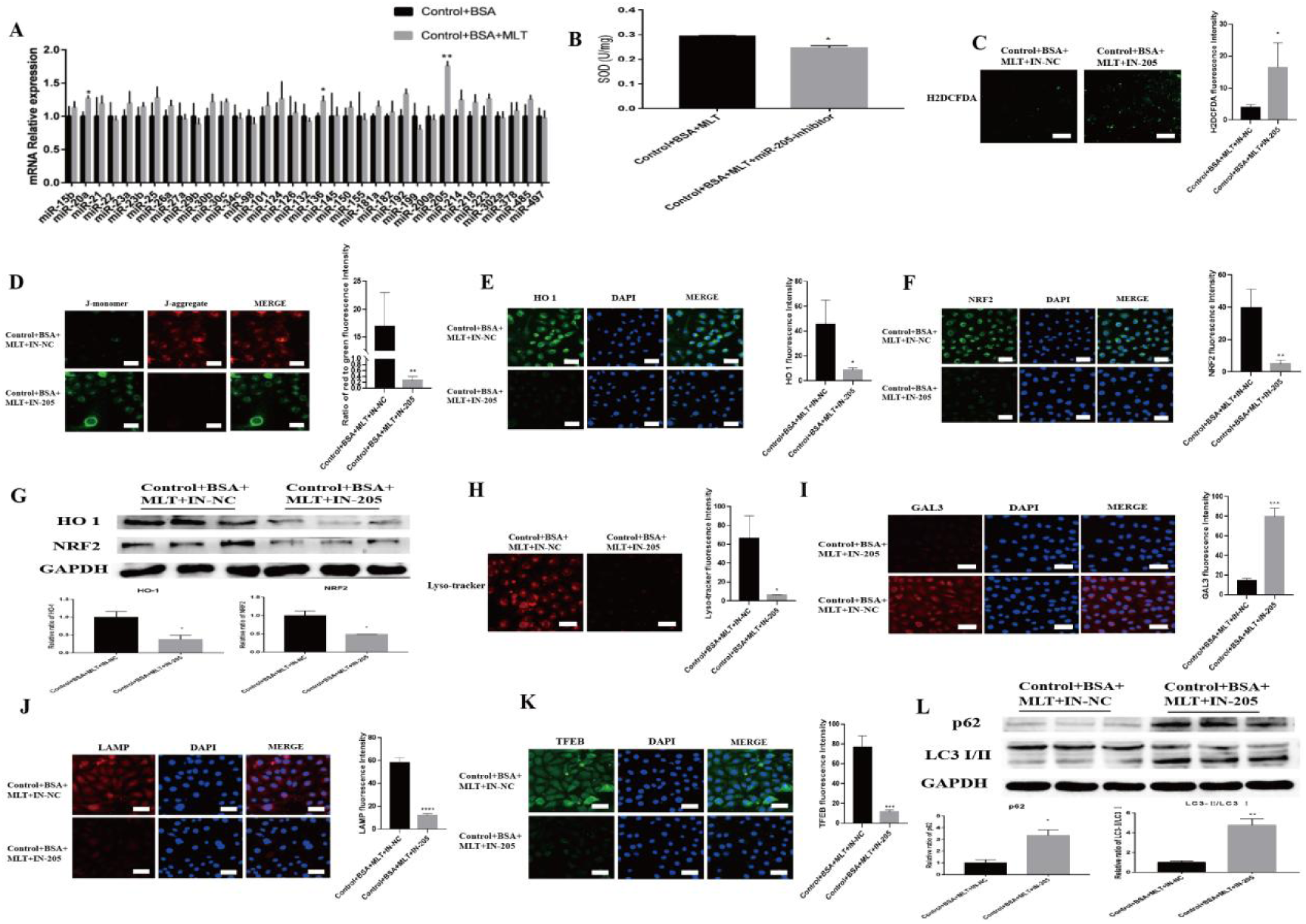
MLT ameliorates lysosomal damage in BSA-HK-2 cells via miR-205-5p (A) Relative miRNA expression after treatment of BSA-HK-2 cells with MLT (n=3/group). ∗ P < 0.05, ∗ ∗ P < 0.01 vs. the control group. (B) SOD activity after miR-205-5p inhibition. ∗ P < 0.05 vs. the control group. (C) Fluorescence intensity of ROS after miR-205-5p inhibition. ∗ P < 0.05 vs. the control group. (D) Images of JC-1 staining after miR-205-5p inhibition. ∗ ∗ P < 0.01 vs. the control group. (E, F) Immunofluorescence staining for HO 1 and NRF2 after miR-205-5p inhibition. ∗ P < 0.05, ∗ ∗ P < 0.01 vs. the control group. (G) Relative HO 1 and NRF2 protein levels after miR-205-5p inhibition. ∗ P < 0.05 vs. the control group. (H) Fluorescence intensity of lysosomal acidification after miR-205-5p inhibition. ∗ P < 0.05 vs. the control group. (I-K) Images of GAL3, LAMP1 and TFEB staining after miR-205-5p inhibition. ∗ ∗ ∗ P < 0.001, ∗ ∗ ∗ ∗ P < 0.0001 vs. the control group. (L) Relative p62 and LC3 protein levels after miR-205-5p inhibition. ∗ P < 0.05, ∗ ∗ P < 0.01 vs. the control group.

### MiR-205-5p downregulates LRP-1 expression to mitigate lysosomal damage in BSA-HK-2 cells

To gain deeper insight into how MLT modulates miR-205-5p to protect against DKD, TargetScan, miRTarBase, miRDB and LGDB were used to predict downstream targets of miR-205-5p. The databases predicted three genes (LRP-1, MMD and NSF) as target genes for miR-205-5p (Figure 4A). The mRNA levels of LRP-1 and MMD were decreased in BSA-HK-2 cells (Figure 4B). After the overexpression of miR-205-5p in HK-2 cells, only the expression of LRP-1 was reduced, while the expression of MMD and NSF did not significantly change (Figure 4C). Next, we predicted the binding site of miR-205-5p and LRP-1 using the TargetScan database (Figure 4D) and validated the results through dual-luciferase reporter gene experiments (Figure 4E). Moreover, in HK-2 cells, miR-205-5p suppressed the protein level of LRP-1 (Figure 4F, G). LRP-1 expression was also significantly decreased in db/db mice overexpressing miR-205-5p (Figure 4H, I), and immunofluorescence staining for LRP-1 was decreased in the miR-205 mimic-treated HK-2 cells (Figure S5). These results indicate that LRP-1 is indeed a direct target regulated by miR-205-5p.

**Figure 4.**
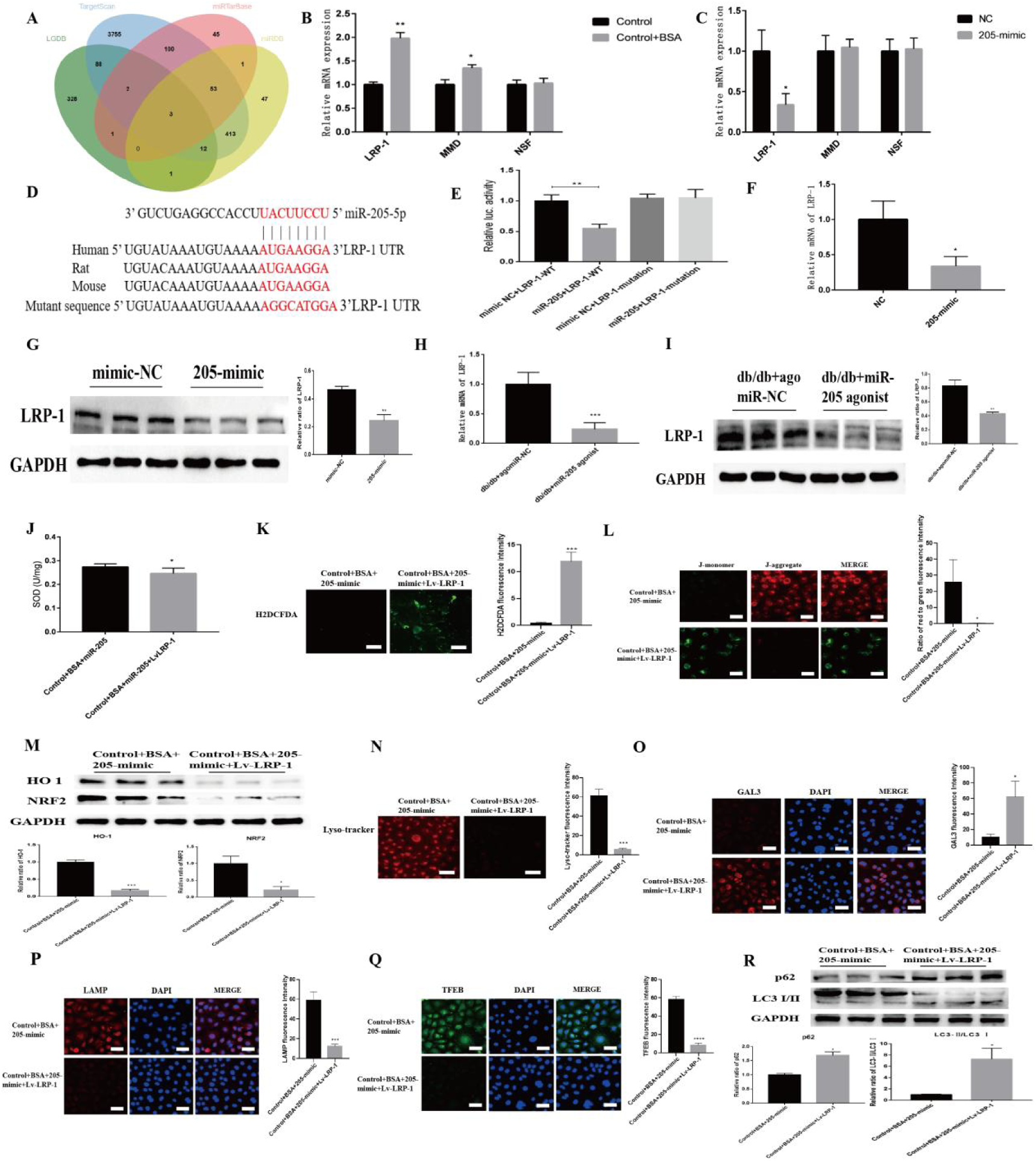
MiR-205-5p ameliorates lysosomal damage in BSA-HK-2 cells via LRP-1 (A) Number of transcripts predicted by miRTarBase, TargetScan, miRDB and LGDB with conserved miR-205-5p binding sites. (B) Relative mRNA expression of LRP-1, MMD and NSF in BSA-HK-2 cells. ∗ P < 0.05, ∗ ∗ P < 0.01 vs. the control group. (C) Relative mRNA expression of LRP-1, MMD and NSF in HK-2 cells overexpressing miR-205-5p. ∗ P < 0.05 vs. the control group. (D) Schematic diagram of the predicted binding sites of miR-205-5p and the mutation in the binding sites of the LRP-1 3’UTR. (E) HEK293T cells were transfected with a dual-luciferase reporter vector containing the wild-type or mutant LRP-1 3’UTR, miR-205-5p mimics or NC mimics, and luciferase activity was measured. ∗ ∗ P < 0.01 vs. the control group. (F) Relative LRP-1 mRNA expression in HK-2 cells. ∗ P < 0.05 vs. the control group. (G) Relative LRP-1 protein expression in HK-2 cells. ∗ ∗ P < 0.01 vs. the control group. (H) Relative LRP-1 mRNA expression in miR-205-overexpressing mice.∗ P < 0.05 vs. the control group. (I) Relative LRP-1 protein expression in miR-205-overexpressing mice. ∗ ∗ P < 0.01 vs. the control group. (J) SOD activity after LRP-1 overexpression. ∗ P < 0.05 vs. the control group. (K) Fluorescence intensity of ROS after LRP-1 overexpression. ∗ ∗ ∗ P < 0.001 vs. the control group. (L) Staining images of JC-1 after LRP-1 overexpression. ∗ P < 0.05 vs. the control group. (M) Relative HO 1 and NRF2 protein levels after LRP-1 overexpression. ∗ P < 0.05 vs. the control group. (N) Fluorescence intensity of lysosomal acidification after LRP-1 overexpression. ∗ ∗ ∗ P < 0.001 vs. the control group. (O-Q) Images of GAL3, LAMP1 and TFEB staining after LRP-1 overexpression. ∗ P < 0.05, ∗ ∗ ∗ P < 0.001, ∗ ∗ ∗ ∗ P < 0.0001 vs. the control group. (R) Relative p62 and LC3 protein levels after LRP-1 overexpression. ∗ P < 0.05 vs. the control group.

To further clarify the mechanism by which miR-205-5p alleviates lysosomal damage, we transfected BSA-HK-2 cells with a miR-205-5p mimic with or without concomitant LRP-1 overexpression (Figure S6A, B). Compared with BSA-HK-2 cells overexpressing only miR-205-5p, those overexpressing LRP-1 had a lower SOD content (Figure 4J) and a higher ROS content (Figure 4K), which resulted in a significant decrease in the mitochondrial membrane potential (Figure 4L). Significant downregulation of NRF2 and HO 1 expression was also observed (Figure 4M, Figure S7A, B) after LRP-1 overexpression. In addition, fluorescence staining revealed significant lysosomal damage and reduced biosynthesis after LRP-1 overexpression (Figure 4N-Q). The protein levels of LC3 and p62 were also significantly increased (Figure 4R), which suggests the occurrence of autophagy arrest after LRP-1 overexpression. Collectively, these findings indicate that miR-205-5p exerts a protective effect on DKD by regulating LRP-1.

### MLT regulates the expression of miR-205-5p through TFEB

Interestingly, when the JASPAR database was used to predict the upstream regulators of miR-205-5p, TFEB, a crucial factor in regulating lysosomal growth, was considered a potential regulatory factor of miR-205-5p. Sequence alignment analysis revealed a binding site between TFEB and the miR-205-5p promoter region (Figure 5A). We subsequently found that TFEB overexpression (Figure S8A, B) reduced the luciferase activity of the WT miR-205-5p promoter but had no significant effect on the corresponding mutant (Figure 5B). Moreover, in HK-2 cells, TFEB increased the mRNA expression of miR-205-5p (Figure 5C). These results suggest that TFEB acts as a transcription factor that enhances miR-205-5p expression.

**Figure 5.**
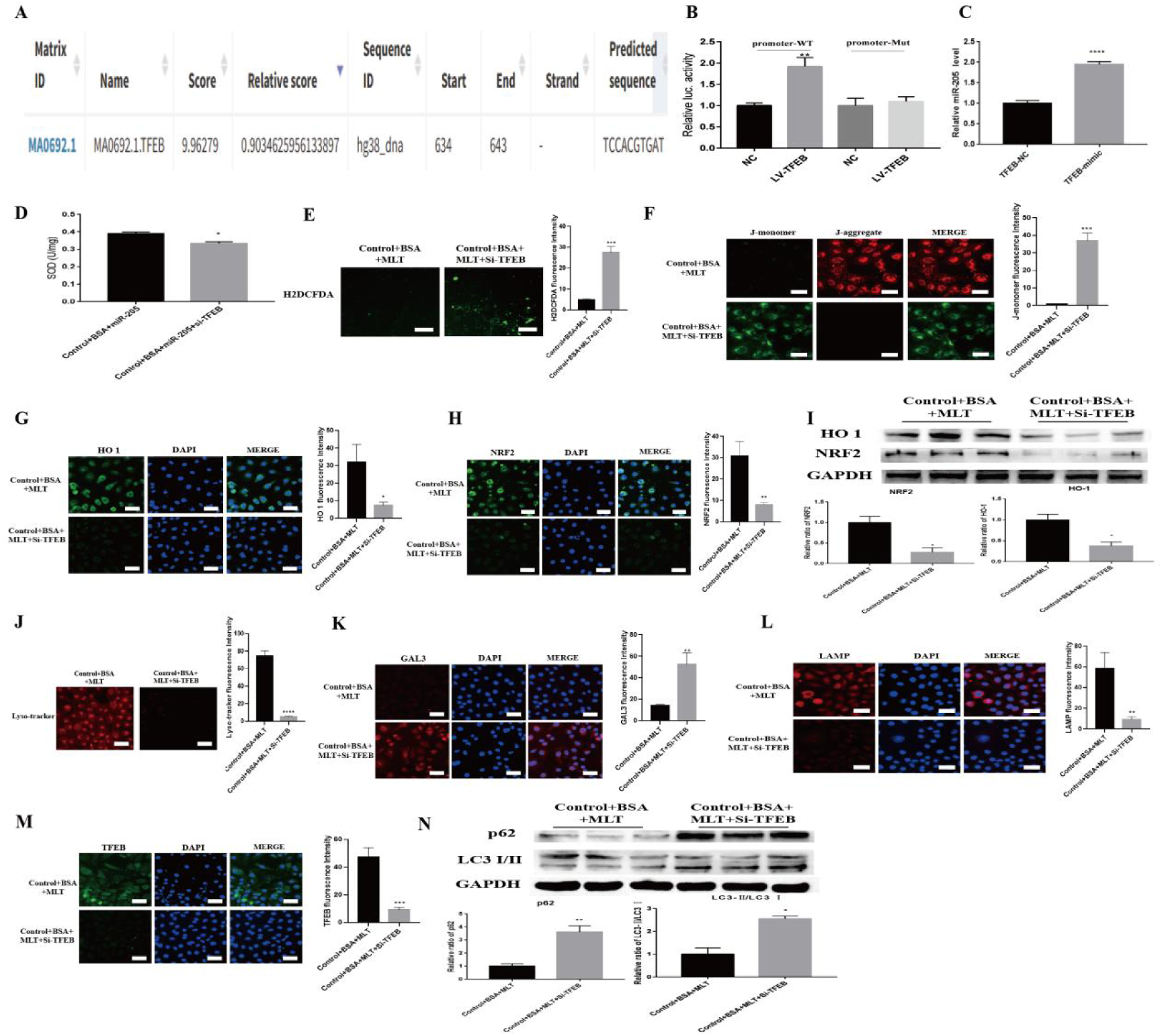
MLT upregulates miR-205-5p through TFEB (A) Schematic diagram of the binding sites of miR-205-5p and TFEB predicted by JASPAR. (B) HEK293T cells were transfected with a dual-luciferase reporter vector containing the WT or mutant miR-205-5p promoter, treated with the TFEB plasmid or NC plasmid, and luciferase activity was measured. ∗ ∗ P < 0.01 vs. the control group. (C) Relative miR-205-5p mRNA expression. ∗ ∗ ∗ ∗ P < 0.0001 vs. the control group. (D) SOD activity after TFEB knockdown. ∗ P < 0.05 vs. the control group. (E) Fluorescence intensity of ROS after TFEB knockdown. ∗ ∗ ∗ P < 0.001 vs. the control group. (F) Staining images of JC-1 after TFEB knockdown. ∗ ∗ ∗ P < 0.001 vs. the control group. (G, H) Immunofluorescence staining for HO 1 and NRF2 after TFEB knockdown. ∗ P < 0.05, ∗ ∗ P < 0.01 vs. the control group. (I) Relative HO 1 and NRF2 protein levels after TFEB knockdown. ∗ P < 0.05 vs. the control group. (J) Fluorescence intensity of lysosomal acidification after TFEB knockdown. ∗ ∗ ∗ ∗ P < 0.0001 vs. the control group. (K-M) Images of GAL3, LAMP1 and TFEB staining after TFEB knockdown. ∗ ∗ P < 0.01, ∗ ∗ ∗ P < 0.001 vs. the control group. (N) Relative p62 and LC3 protein levels after TFEB knockdown. ∗ P < 0.05 vs. the control group. ∗ ∗ P < 0.01 vs. the control group.

To verify whether MLT functions by regulating TFEB and subsequently affecting miR-205-5p, we knocked out TFEB (Figure S9A, B) in BSA-HK-2 cells treated with MLT. Compared with BSA-HK-2 cells treated with MLT alone, SOD activity was lower (Figure 5D) and the ROS level was greater (Figure 5E) in TFEB knockout cells, which resulted in a significant decrease in the mitochondrial membrane potential after TFEB knockdown (Figure 5F). Moreover, immunofluorescence staining (Figure 5G, H) and WB (Figure 5I) revealed that the levels of the antioxidative stress-related proteins NRF2 and HO1 were decreased after TFEB knockdown. In addition, fluorescence staining revealed significant lysosomal damage and reduced biosynthesis after TFEB knockdown (Figure 5J-M). The protein levels of LC3 and p62 were also significantly increased (Figure 5N), which suggests the occurrence of autophagy arrest after TFEB knockdown. These results indicate that TFEB knockdown can block the therapeutic effect of MLT on BSA-HK-2 cells.

### MLT alleviates oxidative stress and lysosomal damage in the renal cortex by upregulating miR-205-5p and downregulating LRP-1 in db/db mice

Compared with db/db mice treated with ddH2O, db/db mice treated with MLT presented increased SOD levels (Figure 6A) and decreased ROS levels (Figure 6B), which increased the mitochondrial membrane potential (Figure 6C). Moreover, immunofluorescence staining (Figure 6D, E) and WB (Figure 6F) revealed that the levels of the antioxidative stress-related proteins NRF2 and HO 1 in db/db mice were significantly increased after treatment with MLT. Moreover, fluorescence staining revealed lysosomal damage repair and increased biosynthesis in db/db mice treated with MLT (Figure 6G-J). The protein expression levels of LC3 and p62 were significantly decreased (Figure 6K), which suggests a decrease in autophagy arrest after MLT treatment.

**Figure 6.**
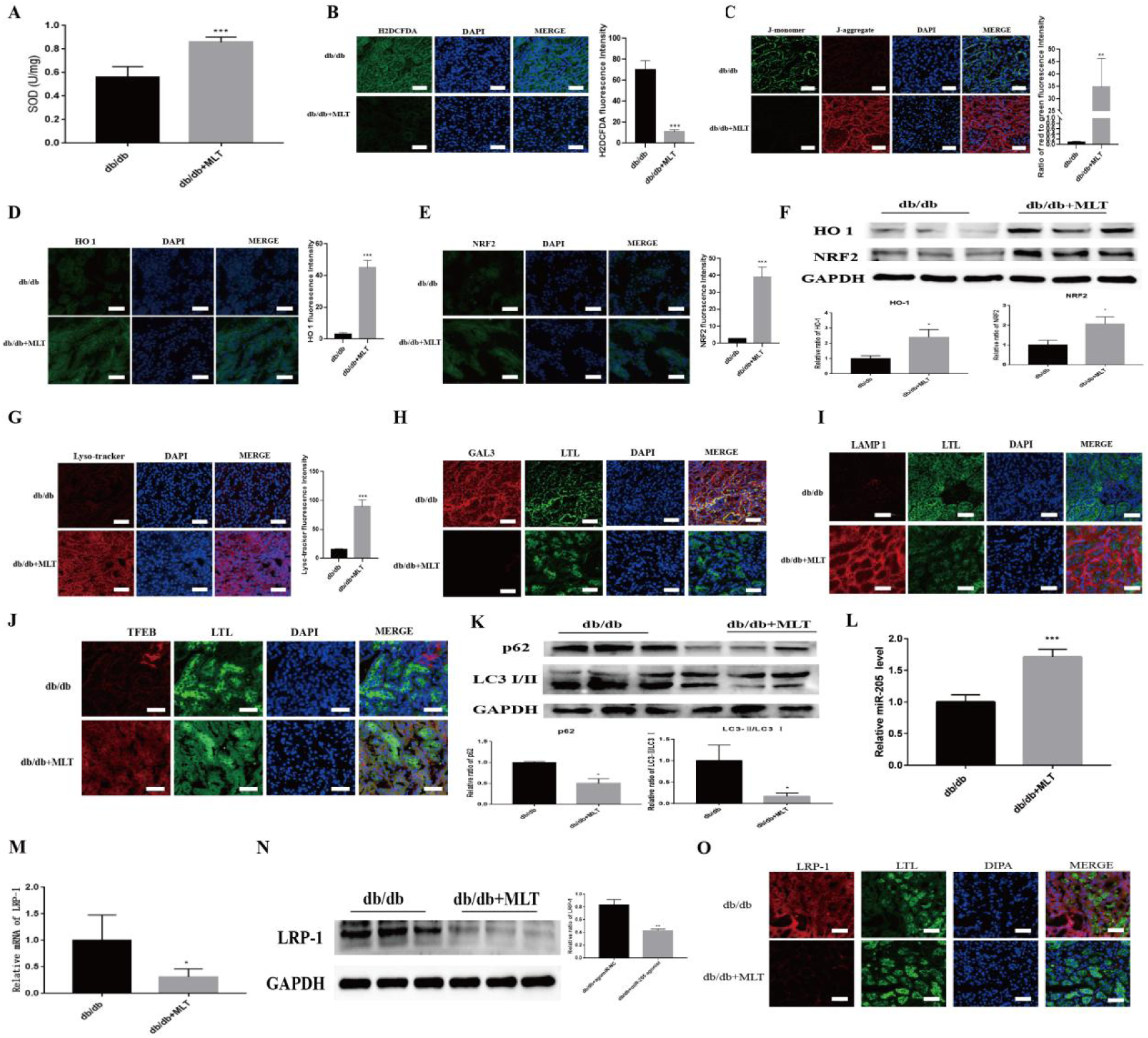
MLT upregulates miR-205-5p and downregulates LRP1 to ameliorate kidney injury in db/db mice (A) SOD activity in MLT-treated db/db and control mice (n=5/group). ∗ ∗ ∗ P < 0.001 vs. the control group. (B) Fluorescence intensity of ROS in MLT-treated db/db and control mice. ∗ ∗ ∗ P < 0.001 vs. the control group. (C) Staining images of JC-1 in MLT-treated db/db and control mice. ∗ ∗ P < 0.01 vs. the control group. (D, E) Immunofluorescence staining for HO 1 and NRF2 in MLT-treated db/db and control mice. ∗ ∗ ∗ P < 0.001 vs. the control group. (F) Relative HO 1 and NRF2 protein levels in MLT-treated db/db and control mice. ∗ P < 0.05 vs. the control group. (G) Fluorescence intensity of lysosomal acidification in MLT-treated db/db and control mice. ∗ ∗ ∗ P < 0.001 vs. the control group. (H-J) Images of GAL3, LAMP1 and TFEB staining in MLT-treated db/db and control mice. (K) Relative p62 and LC3 protein levels in MLT-treated db/db and control mice. ∗ P < 0.05 vs. the control group. (L) Expression of miR-205-5p in MLT-treated db/db and control mice. (n=5/group). ∗ ∗ ∗ P < 0.001 vs. the control group. (M) Relative mRNA expression of LRP-1 in MLT-treated db/db and control mice. ∗ ∗ ∗ P < 0.001 vs. the control group. (N) Relative protein expression of LRP-1 in MLT-treated db/db and control mice. ∗ ∗ ∗ P < 0.001 vs. the control group. (O) Immunofluorescence staining for LRP-1 in MLT-treated db/db and control mice.

**Figure 7.**
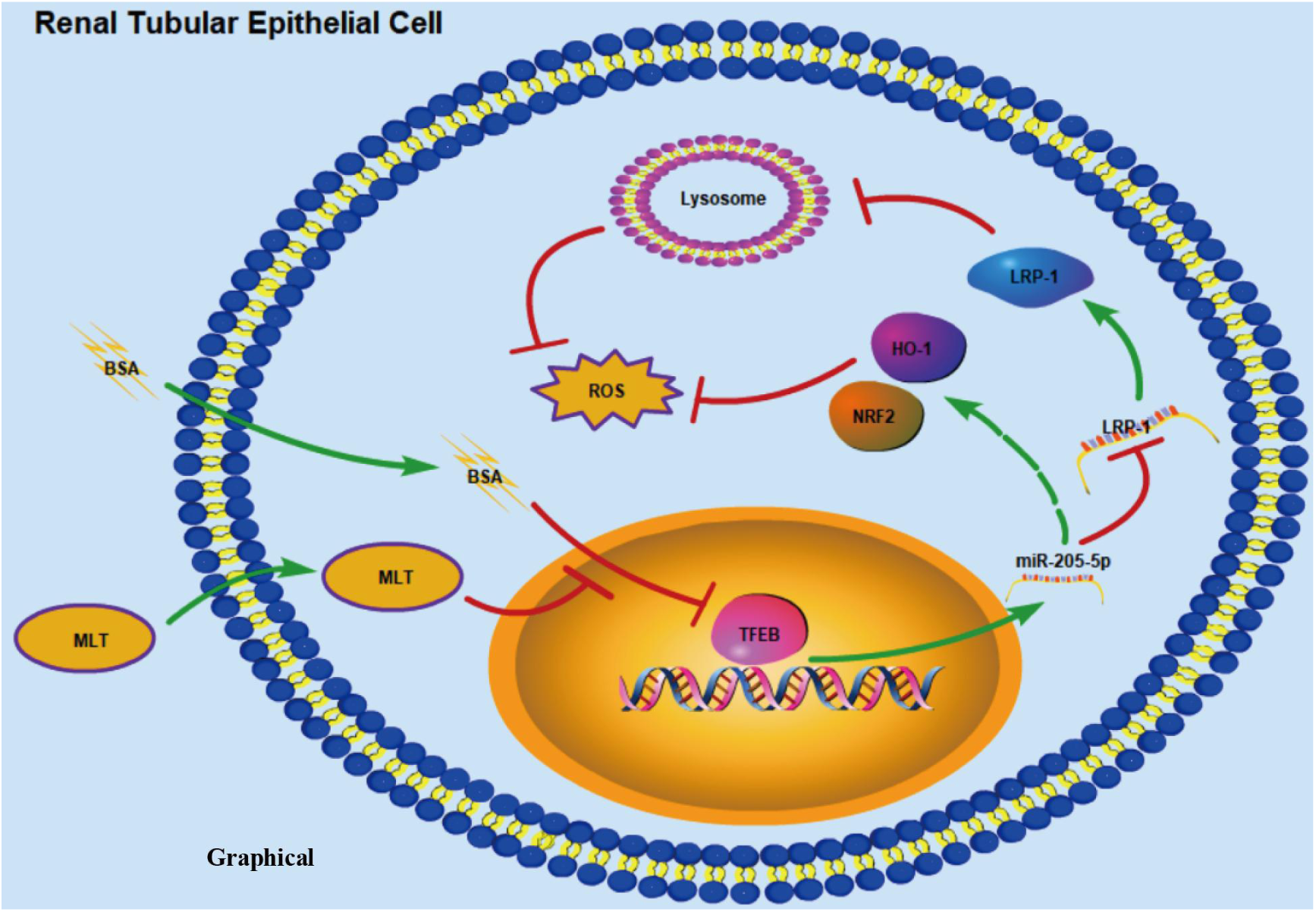
Pattern diagram

Furthermore, in db/db mice treated with MLT, we observed a significant upregulation of miR-205-5p expression (Figure 6L), along with decreased mRNA (Figure 6M) and protein (Figure 6N, O) expression of LRP-1. These results suggest that MLT reduces kidney damage in DKD by increasing miR-205-5p expression and downregulating LRP-1 expression, thereby decreasing oxidative stress and reducing lysosomal damage in the proximal renal tubular region.

## DISCUSSION

Recent research has shown that damage to lysosomes can lead to the instability of renal tubular epithelial cells, which potentially influences DKD development [28]. However, therapeutic interventions for lysosomal damage are still lacking. MLT has been recognized for its role in enhancing autophagy, combating oxidative stress, and addressing other related issues [17, 29]. As a result, MLT has considerable potential for repairing lysosomal damage. In this study, we demonstrated that MLT treatment significantly ameliorated oxidative stress and lysosomal damage in renal tubular epithelial cells, thereby delaying the development of DKD. We found that MLT drives this process by regulating the TFEB-miR-205-LRP-1-lysosomal pathway.

The pathogenesis of DKD is related to multiple factors, including inflammatory cascade reactions, oxidative imbalance, albumin overload, and metabolic disorders such as persistent hyperglycemia [30]. Increasing evidence indicates that impaired autophagy leads to impaired clearance of abnormal organelles and proteins, which in turn disrupts cellular homeostasis and promotes the occurrence of DKD [31]. However, the role and intrinsic mechanisms of lysosomes, which constitute the core component of autophagy, have not been fully elucidated. Liu Weijing and colleagues noted that significant lysosomal damage is present in the kidneys of patients with DKD and that the alleviation of lysosomal injury can significantly delay the occurrence of renal fibrosis in end-stage DKD [16]. However, the current means to improve lysosomal damage are very limited. In this study, we observed that diabetic mice and a cell model presented obvious lysosomal damage and that alleviating lysosomal damage can restore autophagic flux, decrease oxidative stress, and delay the development of DKD.

Previous studies have shown that MLT, the main hormone secreted by the pineal gland, can delay the progression of DKD by alleviating kidney inflammation and oxidative stress in DKD models [32, 33]. A clinical trial of MLT supplements in diabetic patients demonstrated that MLT administration may be a new treatment to improve diabetic status and reduce the incidence of diabetic complications [34]. In further mechanistic studies, MLT was also found to ameliorate endothelial damage to glomerular podocytes and endothelial cells in in vitro models, in which MLT significantly alleviates DKD. Tubular injury is one of the "culprits" of DKD, but the mechanism by which MLT treatment exerts protective effects has not been determined. Our study revealed that MLT alleviated disturbances in autophagy and oxidative stress in the kidneys of diabetic mice and in cells stimulated with albumin in an in vitro diabetic kidney disease model. Several previous studies reported that MLT can play an important protective role by alleviating lysosomal injury caused by neurological and cardiovascular diseases [35, 36]. Thus, lysosomes may also be promising targets through which MLT can ameliorate renal tubular damage. Our results also revealed that MLT restored autophagy pathway activity and antioxidation resistance by decreasing lysosomal damage.

Recent studies have strongly linked miRNAs with the development of DKD and suggest that they may be valuable targets for innovative therapeutic approaches [37, 38]. Our study revealed that the expression of renal-protective miR-205-5P significantly increased after treatment with MLT, a finding that is consistent with the results of our previous study. To identify downstream targets, we compared the proteins obtained by miR-205-5P downstream target prediction with the lysosomal proteins associated with miR-205-5p and selected LRP-1 as a downstream target candidate. LRP-1 is a transmembrane protein belonging to the LDL receptor family that is associated with lipoprotein metabolism and cellular homeostasis [39]; LRP-1 plays a key regulatory role in various cellular activities, including proliferation, motility, differentiation, and transdifferentiation, and the aberrant expression of LRP-1 can cause organ dysfunction and damage [40]. Notably, LRP-1 expression is significantly elevated during kidney injury, and this elevation can activate profibrosis-related signaling pathways, which subsequently aggravate fibrosis [41, 42]. Moreover, studies have reported that high LRP-1 expression can promote lysosomal degradation through ligands such as PCSK9 [43]. Our study revealed that MLT could upregulate miR-205-5P expression, downregulate LRP-1 expression, alleviate lysosomal damage, restore autophagic flux, resist oxidative stress, and ultimately delay the development of DKD.

Interestingly, when we explored how MLT regulates the expression of miR-205-5p, we discovered that transcription factor EB (TFEB), which is involved in lysosomal biogenesis, can upregulate miR-205-5p. Moreover, we also observed an increase in TFEB protein levels and nuclear entry after MLT treatment, which indicates that MLT may restore autophagic flux by increasing TFEB expression and activation. We subsequently verified that TFEB could regulate the expression of miR-205-5p. These results indicate that MLT can not only ameliorate lysosomal damage and oxidative stress through the TFEB-miR-205-LRP1 pathway but can also directly regulate TFEB to promote lysosome biogenesis and subsequently improve the autophagy‒lysosome pathway. These findings provide strong evidence for the therapeutic potential of MLT for patients with DKD.

Collectively, our findings indicate that MLT improves lysosomal damage in renal tubular epithelial cells through TFEB and its downstream pathways, such as the miR-205‒LRP-1 pathway, and ultimately contributes to the delayed progression of DKD. Importantly, MLT has enormous potential as a drug for the treatment of DKD.

## Acknowledgments and GRANTS

Our study was supported by grants from the National Natural Science Foundation of China (Z.Z., 82270862, 81700730) (Y.J., 82370818, 82000785); the Guangzhou Science and Technology Project (Y.J., 2024A04J5098); the National Undergraduate Training Program for Innovation and Entrepreneurship, Southern Medical University (Z.Z., 202312121031, Y.J., S202312121167); and the Bethune Charitable Foundation (B-0307-H- 20200302).

## AUTHOR CONTRIBUTIONS

Z.Z., Y.J., Y.X., and J.C. conceived and designed the research; J.C., S.Z., X.M., A.C., Y.W. and Z.Z. performed the experiments; J.C., X.X., S.Z., X.M., A.C., Y.W. and G.W. analyzed the data; J.C., S.Z., interpreted the results of the experiments; J.C., and S.Z., prepared the figures; J.C., and S.Z., drafted the manuscript; Z.Z., Y.J., Y.X., and J.C. edited and revised the manuscript; all the authors made substantial contributions to the manuscript and approved it for submission. Z.Z. and Y.J. and Y.X. are the guarantors of this work, and as such, had full access to all the data in the study and take responsibility for the integrity of the data and the accuracy of the data analysis.

