## Supplementary Materials for "Melatonin attenuates kidney injury by alleviating lysosomal damage in diabetic kidney disease"

### Supplementary Tables

#### Table S1. Primers used for quantitative RT-PCR.

| Host | Gene | Primer sequence (5’-3’) |
|---|---|---|
| Human | LRP-1 | F: GGCTGATCAGGTGTCGGAAA |
| | | R: GTCACCGTTGTTGACACTGC |
| | MMD | F: ATGGAATGGGACTCTGTGCC |
| | | R: CAGCTGCCATGAGCCAGATA |
| | NSF | F: TGCTGGTGCAGCAGACTAAG |
| | | R: TCATGGCCTGACATTTGGCT |
| | TFEB | F: TCTGTCCAGCAACATGACAGC |
| | | R: GGATGTGGGATTCTCCAGGT |
| Mouse | LRP-1 | F: GCGGTGTGACAACGACAATG |
| | | R: GGTCTTGTAGCCTGGTTGGT |
| | | R: TCTATCGGATACTTCAGCGTCA |

#### Table S2. The sequence of wild-type or mutant LRP-1 3'-UTR

| | |
|---|---|
| Plasmid | h- LRP-1 ENST00000243077.3 3' UTR-WT |
| Sequence | ctcgagGgccctgccccgtcggactgcccccagaaagcctcctgccccctgccagtgaagtccttcagtgagc<br>ccctccccagccagcccttccctggccccgccggatgtataaatgtaaaaatgaaggaattacattttatatgtgag<br>cgagcaagccggcaagcgagcacagtattatttctccatcccctccctgcctgctccttggcacccccatgctgcc<br>ttcagggagacaggcagggagggcttggggctgcacctcctaccctcccaccagaacgcaccccactgggag<br>agctggtggtgcagccttcccctccctgtataagacactttgccaaggctctcccctctcgccccatccctgcttgc<br>ccgctcccacagcttcctgagggctgcggccgc |
| Plasmid | h- LRP-1 ENST00000243077.3 3' UTR-MUT |
| Sequence | ctcgagGgccctgccccgtcggactgcccccagaaagcctcctgccccctgccagtgaagtccttcagtgagc |

| | |
|---|---|
| | ccctccccagccagcccttccctggccccgccggatgtataaatgtaaaaaggcatggattacattttatatgtgag<br>cgagcaagccggcaagcgagcacagtattatttctccatcccctccctgcctgctccttggcacccccatgctgcc<br>ttcagggagacaggcagggagggcttggggctgcacctcctaccctcccaccagaacgcaccccactgggag<br>agctggtggtgcagccttcccctccctgtataagacactttgccaaggctctcccctctcgccccatccctgcttgc<br>ccgctcccacagcttcctgagggctgcggccgc |

**Table S3. The sequence of wild-type or mutant miR-205-5p promoter**

| | |
|---|---|
| Plasmid | h-miR-205-5p promoter hg38_dna range=chr1:209426820-209428820 promoter-WT |
| Sequence | gctagcGTGAATGAAATCCACAGGCCCATCACAAGATGGCTGCGCTCT<br>CCAGTCCCCGGCACCCCAGGCTGCTCTGAACATGTGAGAGCTCCTCC<br>CTCAAGGCCTGAATTTTCGCAGGTCTTCATCACACTCAGCCCGCCAT<br>CATTTTTTCAGATTCAGTTCCAATATGTCACTCTGTCTGTAGAGCCTT<br>CCCTGAATCTCACAGACAAATGCGGCAGTTCCCTTTGTGGGGGATAT<br>TCATCATGTTTTTCCTCATTGGCCTATGTTTTTGCCTCTTCTTGTAGAT<br>GTCCATCTAGGCTGGGAGCTCCTAGGGGGTAAACGCAGCCTCTCAC<br>TCACCTGGCACAGAGTCGATGCTCATGGAATGATACGTGATGGGGA<br>GAGTGAGTAAGTGAATGAATGAATGAATGCTATTCTGGAAGGAAGA<br>GGGAGAGAGGGAGCAGGGAAAGGTCAGAAGGGGGCCCAGAGTCTT<br>CAGCACACAATGTGGGTGTATCACTGTGGCCAGGAACAATTCGCTT<br>CTCTGCCTGCATTACTTTTTGGGAAATGGGAGAGCTGAGGCTCAGTG<br>TAGGCAAACTGGGTCCAGTCAGCAGGGAAGCCATGAGCTCCACGGT<br>TTAGGGGAGGAGTCCGGTGCTTAAGGAAGCTATCACGTGGAAGATC<br>TCCCTGCCCTCCTCCTATTGGTCTGTATTGGTTTTTCTTCTTCACCAG<br>TGGAGGCTTTTCTTCTCCCAGGAAGAATCCAGAATAGTTCAGACAA<br>GCTTCAGGTCCGCCAGAACCGGGGAAAACAACGTGTAGGGTTTGTT<br>TTAAAGGGCTTTTAAATGGGGTTTGGGAGATCCAGGTAGATTAGAA<br>TCTTGACTCTGCACTGGCTGTGCAAGACAGAAATTCACCTTTCAAAA |

| | |
|---|---|
| | CTTCTGTTCCCCAGTCTATGCGGTAAGAAGTTTAGATTAGCTGATTT<br>CTAGGGAGATTTTGGGCAGGCATGATTCAGTTCCATGAAGGGCTGT<br>GGGACAGGACTGCACTAAGAGGTCCTATAGATTGGATTCTTCAGTTC<br>CCTCCCTTCCGCTGGAGAATAGAGGAATCTGCCTTCGCTGCACACTA<br>GATTTACCTGGAGAACAAATGGAAAGCAGGCCAGGATCCTGGGTGA<br>AATGTGGCACATTTGAGGAGACTTCAACTTCCCTTGCCCTCCAGGTG<br>TGCACTTGGAAGGGGGAACGAGGGAGGGGGAAGCTGGCAAGATGG<br>GCCGAGAGATAGGGGAGGGGCAGGAGGCGGAGCCCAAGTTGCGCA<br>TGGAAGCGGGGGTGGGGTAAAATAATCAAGTTTATAGACCGCCCTC<br>TTTAAAGTTACTAATGAGCTTGCCTTCTCTTTCCTTAATTTCCCCTCG<br>CAGTGTGGTCTTTTCCCCACCCCCAGACATGAAAGGGAAGCAGGTC<br>ACAAAGCCTTTCGGATTATAAAAGAAACACTTGCTTCTCACAAGGG<br>GAGCAGCAGACTTACTCTGTACTAAATGCCAGGATAAGCCTCTGGC<br>TGGGCCTCGACTGTGACCCTCCGGCCTCTTTCTACAGCTCTGCCTGG<br>ATGGACTGGCCTATCTCTGCTGGATTCCCGAAGTGCATTGTGTAGAG<br>ACAGCAACTCAGGTCAGGCTAAAAGCTCAAGCAAGCAAGCGCGCAC<br>ACACACGCGCGCACACACACACACACACACACAAACACTCAGCTTC<br>CTTAGGACAAGATAAAATCTTAGCATTCCCCTCTCCCCGATTAGGTA<br>GGTCTCTGGGAGGACTAAGGCTTCAGGTGCAAGGCTCAGATAACCT<br>GCAGTGTCTCTCCAACTCTGGGATGACAAAGACCTCACTTCCCTTTT<br>CTGGTGTTCACACAACAATGAGAAAGTACGGGTAGCCTGCAGAAAG<br>ACCTCTCCATTCATGGTCCCCCAGGGGTGTGGGTTCTGAGAGGTGGG<br>ACCAGCTGCCAGGCCCTTTCTCCATTGGTTGAGTTCAGCAGGTAACC<br>TGAAGCTTTGCTGAGAGGTGCATAAATAAAGAGTGAAACTAGTACC<br>ACCTCCgtcgac |
| Plasmid | h-miR-205-5p promoter hg38_dna range=chr1:209426820-209428820<br>promoter-MUT |
| Sequence | gctagcGTGAATGAAATCCACAGGCCCATCACAAGATGGCTGCGCTCT<br>CCAGTCCCCGGCACCCCAGGCTGCTCTGAACATGTGAGAGCTCCTCC |

CTCAAGGCCTGAATTTTCGCAGGTCTTCATCACACTCAGCCCGCCAT
CATTTTTTCAGATTCAGTTCCAATATGTCACTCTGTCTGTAGAGCCTT
CCCTGAATCTCACAGACAAATGCGGCAGTTCCCTTTGTGGGGGATAT
TCATCATGTTTTTCCTCATTGGCCTATGTTTTTGCCTCTTCTTGTAGAT
GTCCATCTAGGCTGGGAGCTCCTAGGGGGTAAACGCAGCCTCTCAC
TCACCTGGCACAGAGTCGATGCTCATGGAATGATACGTGATGGGGA
GAGTGAGTAAGTGAATGAATGAATGAATGCTATTCTGGAAGGAAGA
GGGAGAGAGGGAGCAGGGAAAGGTCAGAAGGGGGCCCAGAGTCTT
CAGCACACAATGTGGGTGTATCACTGTGGCCAGGAACAATTCGCTT
CTCTGCCTGCATTACTTTTTGGGAAATGGGAGAGCTGAGGCTCAGTG
TAGGCAAACTGGGTCCAGTCAGCAGGGAAGCCATGAGCTCCACGGT
TTAGGGGAGGAGTCCGGTGCTTAAGGAAGCTTaCgCtTaAcAGATCTC
CCTGCCCTCCTCCTATTGGTCTGTATTGGTTTTTCTTCTTCACCAGTG
GAGGCTTTTCTTCTCCCAGGAAGAATCCAGAATAGTTCAGACAAGCT
TCAGGTCCGCCAGAACCGGGGAAAACAACGTGTAGGGTTTGTTTTA
AAGGGCTTTTAAATGGGGTTTGGGAGATCCAGGTAGATTAGAATCT
TGACTCTGCACTGGCTGTGCAAGACAGAAATTCACCTTTCAAAACTT
CTGTTCCCCAGTCTATGCGGTAAGAAGTTTAGATTAGCTGATTTCTA
GGGAGATTTTGGGCAGGCATGATTCAGTTCCATGAAGGGCTGTGGG
ACAGGACTGCACTAAGAGGTCCTATAGATTGGATTCTTCAGTTCCCT
CCCTTCCGCTGGAGAATAGAGGAATCTGCCTTCGCTGCACACTAGAT
TTACCTGGAGAACAAATGGAAAGCAGGCCAGGATCCTGGGTGAAAT
GTGGCACATTTGAGGAGACTTCAACTTCCCTTGCCCTCCAGGTGTGC
ACTTGGAAGGGGGAACGAGGGAGGGGGAAGCTGGCAAGATGGGCC
GAGAGATAGGGGAGGGGCAGGAGGCGGAGCCCAAGTTGCGCATGG
AAGCGGGGGTGGGGTAAAATAATCAAGTTTATAGACCGCCCTCTTT
AAAGTTACTAATGAGCTTGCCTTCTCTTTCCTTAATTTCCCCTCGCAG
TGTGGTCTTTTCCCCACCCCCAGACATGAAAGGGAAGCAGGTCACA
AAGCCTTTCGGATTATAAAAGAAACACTTGCTTCTCACAAGGGGAG

CAGCAGACTTACTCTGTACTAAATGCCAGGATAAGCCTCTGGCTGG
GCCTCGACTGTGACCCTCCGGCCTCTTTCTACAGCTCTGCCTGGATG
GACTGGCCTATCTCTGCTGGATTCCCGAAGTGCATTGTGTAGAGACA
GCAACTCAGGTCAGGCTAAAAGCTCAAGCAAGCAAGCGCGCACACA
CACGCGCGCACACACACACACACACACACAAACACTCAGCTTCCTT
AGGACAAGATAAAATCTTAGCATTCCCCTCTCCCCGATTAGGTAGGT
CTCTGGGAGGACTAAGGCTTCAGGTGCAAGGCTCAGATAACCTGCA
GTGTCTCTCCAACTCTGGGATGACAAAGACCTCACTTCCCTTTTCTG
GTGTTCACACAACAATGAGAAAGTACGGGTAGCCTGCAGAAAGACC
TCTCCATTCATGGTCCCCCAGGGGTGTGGGTTCTGAGAGGTGGGACC
AGCTGCCAGGCCCTTTCTCCATTGGTTGAGTTCAGCAGGTAACCTGA
AGCTTTGCTGAGAGGTGCATAAATAAAGAGTGAAACTAGTACCACC
TCCgtcgac

### Supplementary Figures

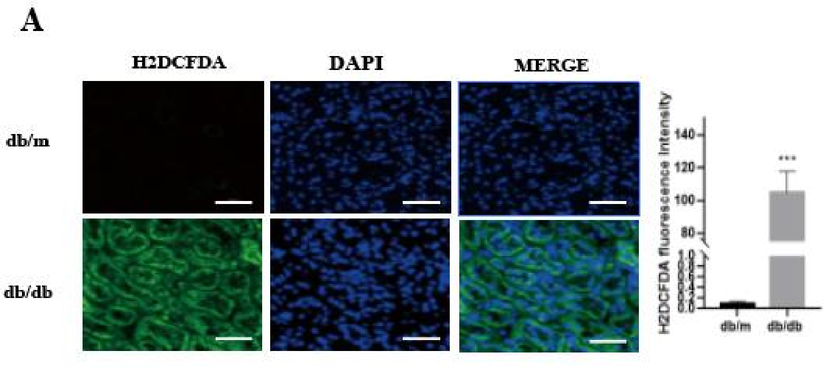


**Figure S1** Fluorescence intensity of ROS in db/db and db/m mice. ＊＊＊ P < 0.001 vs. the db/m group (n=5/group).

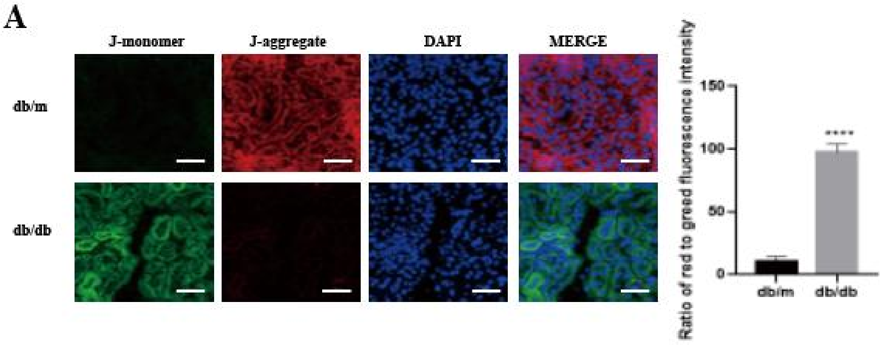


**Figure S2** JC-1 staining images in db/db and db/m mice. ＊＊＊＊ $P < 0.0001$ vs. the db/m group (n=5/group).

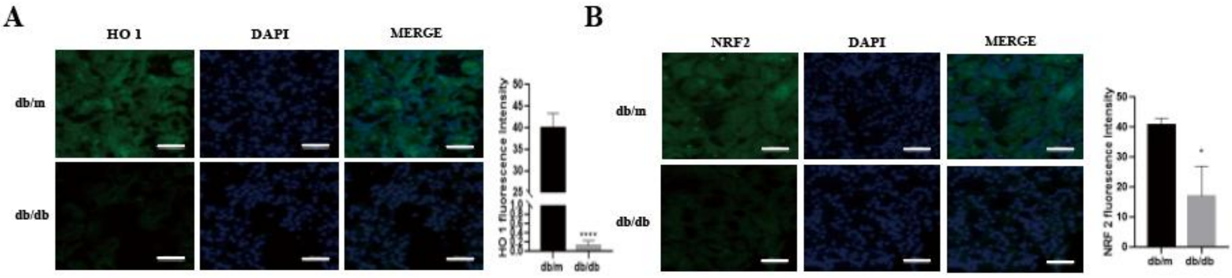


**Figure S3** Immunofluorescence staining for HO 1 and NRF2 in db/db and db/m mice.

(A) Immunofluorescence staining for HO 1 in db/db and db/m mice. ＊＊＊＊ $P < 0.0001$ vs. the db/m group (n=5/group). (b) Immunofluorescence staining for NRF2 in db/db and db/m mice. ＊ $P < 0.05$ vs. the db/m group (n=5/group).

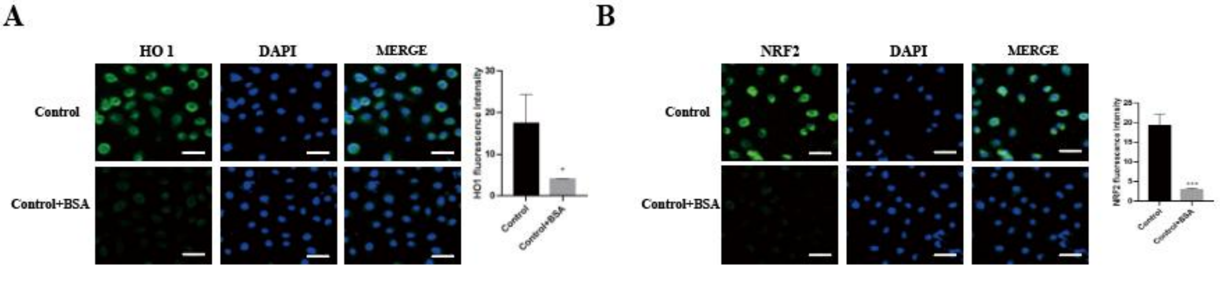


**Figure S4** Immunofluorescence staining for HO 1 and NRF2 in BSA-HK-2 cells and control cells.

(A) Immunofluorescence staining for HO 1 in BSA-HK-2 cells and control cells. ＊ $P < 0.05$ vs. the control group (n=3/group). (B) Immunofluorescence staining for NRF2 in BSA-HK-2 cells and control cells. ＊＊＊ $P < 0.001$ vs. the control group (n=3/group).

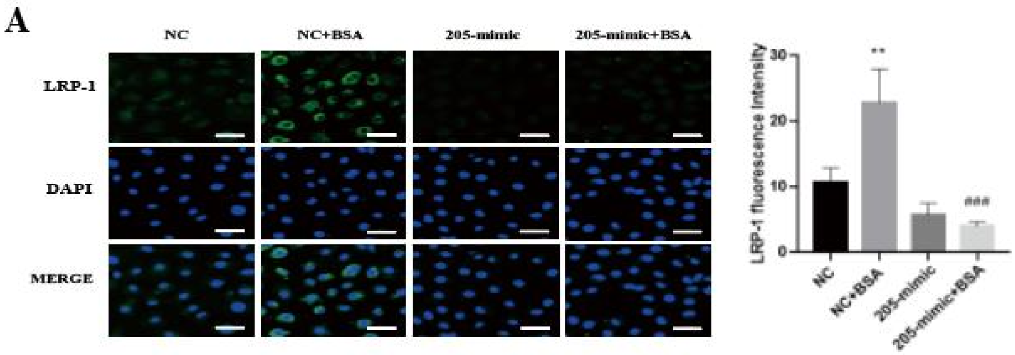


**Figure S5** Immunofluorescence staining for LRP-1 in miR-205 mimic-treated HK-2 cells. ＊＊ P < 0.01 vs. the NC group. ###P < 0.001 vs. the miR-205 mimic group (n=5/group).

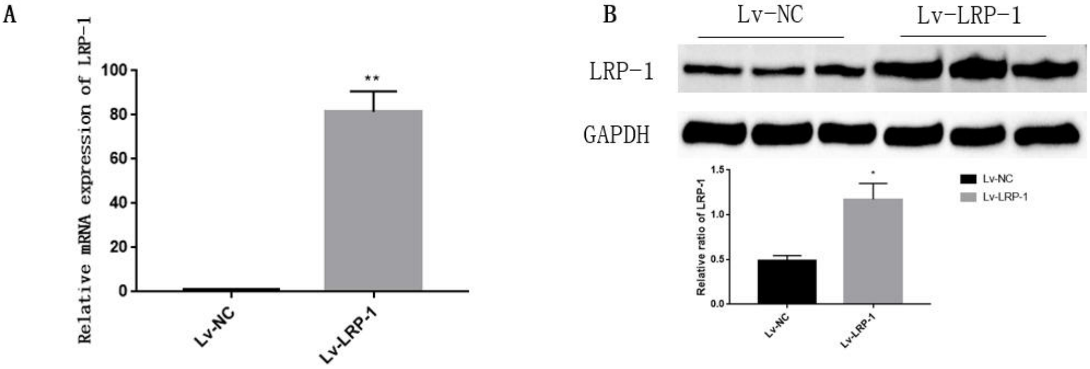


**Figure S6** LRP-1 mRNA and protein expression in HK-2 cells transfected with the pEZ-Lv201-LRP-1 plasmid.

(A) Relative LRP-1 mRNA expression in HK-2 cells transfected with the pEZ-Lv201-LRP-1 plasmid. **P<0.01 compared with the Lv-NC group (n=3/group). (B) Relative LRP-1 protein levels in HK-2 cells transfected with the pEZ-Lv201-LRP-1 plasmid. *P<0.05 compared with the Lv-NC group (n=3/ group).

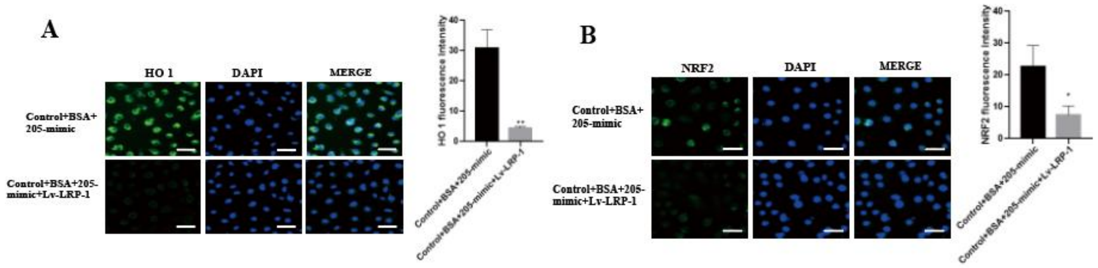


**Figure S7** Immunofluorescence staining for HO 1 and NRF2 in BSA-HK-2 cells transfected with a miR-205-5p mimic with or without concomitant LRP-1 overexpression.

(A) Immunofluorescence staining for HO 1 in BSA-HK-2 cells transfected with a miR-205-5p mimic with or without concomitant LRP-1 overexpression. ＊＊ P < 0.01 vs. BSA-HK-2 cells transfected with a miR-205-5p mimic without concomitant LRP-1 overexpression (n=3/group). (B) Immunofluorescence staining for NRF2 in BSA-HK-2 cells transfected with a miR-205-5p mimic with or without concomitant LRP-1 overexpression. ＊ P < 0.05 vs. BSA-HK-2 cells transfected with a miR-205-5p mimic without concomitant LRP-1 overexpression (n=3/ group).

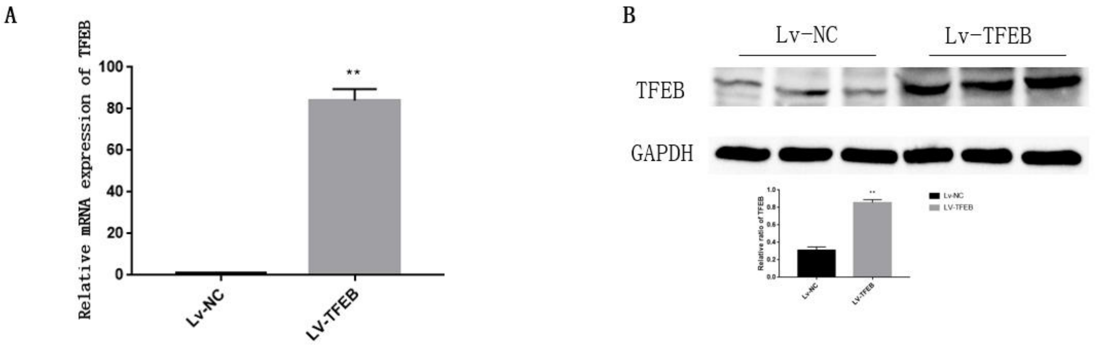


**Figure S8** TFEB mRNA and protein expression in HK-2 cells transfected with the pEZ-Lv201-TFEB plasmid.

(A) Relative TFEB mRNA expression in HK-2 cells transfected with pEZ-Lv201-TFEB plasmid. **P<0.01 compared with the si-NC group (n=3/ group). (B) Relative TFEB protein levels in HK-2 cells transfected with pEZ-Lv201-TFEB plasmid. **P<0.01 compared with the si-NC group (n=3/group).

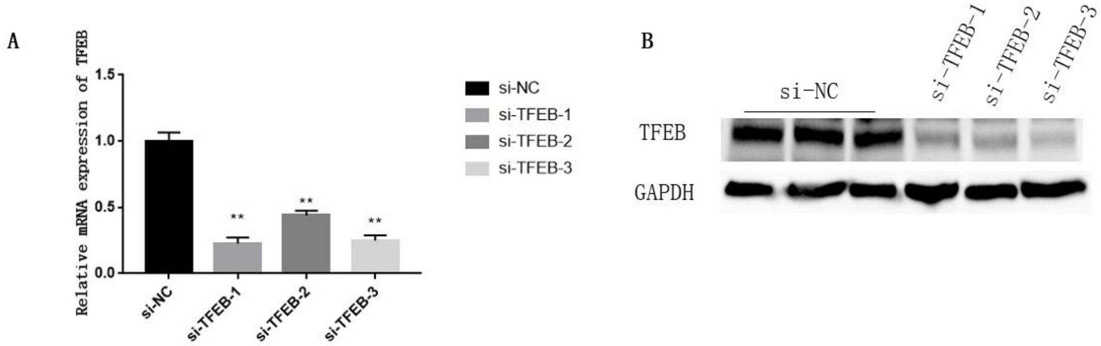


**Figure S9** TFEB mRNA and protein expression in HK-2 cells transfected with si-TFEB.

(A) Relative TFEB mRNA expression in HK-2 cells transfected with si-TFEB. **P<0.01 compared with the si-NC group (n=3/group). (B) Relative TFEB protein levels in HK-2 cells transfected with si-TFEB.
